# Genomic Insights into the Diversity, Antimicrobial Resistance, and Zoonotic Potential of *Campylobacter fetus* Across Diverse Hosts and Geographies

**DOI:** 10.1101/2025.02.16.638542

**Authors:** Ellis Kobina Paintsil, Cynthia Kyerewaa Adu-Asiamah, Kennedy Gyau Boahen, Charity Wiafe Akenten, Alexander Kwarteng, Stefan Berg, Kwasi Obiri-Danso, Jürgen May, Denise Dekker, Linda Aurelia Ofori

## Abstract

**Background:** *Campylobacter fetus* causes reproductive diseases in livestock and zoonotic infections in humans, especially in immunocompromised individuals. Despite its significance, its genomic characteristics are poorly understood. This study analyzed 114 publicly available *C. fetus* genomes to provide global insights into its genetic diversity, antimicrobial resistance (AMR) patterns, and zoonotic risk.

**Results:** A total of 32 distinct sequence types (STs), ranging from ST-1 to ST-74, were identified across 111 of the 114 *C. fetus* genomes, spanning six continents and diverse hosts (cattle, humans, sheep, and reptiles). ST-4 was the most prevalent (n = 45), followed by ST-3 (n = 8). A significant proportion (90.9%; n/N=40/44) of *C. fetus* subsp. *venerealis* (Cfv) and its biovar intermedius (Cfvi) were assigned to ST-4. Despite being isolated from five continents, Cfv and Cfvi genomes clustered closely, forming distinct branches at the biovar level; however, six Cfv genomes were located within Cfvi clades, suggesting a shared evolutionary lineage. In contrast, *C. fetus* subsp. *testudinum* (Cft) genomes, exhibiting 20 distinct STs, formed distinct clades from Cfv, Cfvi, and *C. fetus* subsp. *fetus* (Cff). While Cfv genomes from North America and Cfvi genomes from South America formed distinct geographic clusters, Cff genomes displayed no clear geographical patterns, with branches containing strains from multiple continents, indicating a globally dispersed distribution. Pangenomic analysis revealed pronounced clustering within Cft, characterized by unique gene presence/absence patterns. Five distinct AMR genes were detected, with *tet(O)* (n = 3) being the most common. Horizontal gene transfer analysis identified 140 genomic islands across 41 genomes, and virulence factor analysis revealed *cheY* as the sole conserved virulence gene across 35 genomes.

**Conclusion:** These findings provide critical insights into the genomic diversity, zoonotic potential, and global distribution of *C. fetus*, emphasizing the need for integrated genomic and epidemiological strategies to assess its impact on human and animal health.

## Background

*Campylobacter fetus* is a bacterial species comprising three recognized subspecies: *Campylobacter fetus* subsp. *fetus* (Cff), *Campylobacter fetus* subsp. *venerealis* (Cfv), and *Campylobacter fetus* subsp. *testudinum* (Cft). These subspecies are primarily distinguished by host specificity, ecological niche, pathogenicity, and genetic features (van Bergen et al., 2008). Cff has a broad host range, including sheep, cattle, and humans, where it causes systemic infections and reproductive disorders, such as ovine abortion and bacteremia in immunocompromised individuals (Dumic et al., 2017; Fujihara et al., 2006; Kaakoush et al., 2015). Cfv is host-restricted to cattle, colonizing the reproductive tract and causing bovine genital campylobacteriosis (BGC), a major concern in livestock due to its impact on fertility and early embryonic loss (Sahin et al., 2017; Zan Bar et al., 2008). Cft, primarily associated with reptiles, is an emerging zoonotic pathogen occasionally implicated in human infections, especially in individuals with close reptile contact ((Gilbert et al., 2018; Giacomelli & Piccirillo, 2014). Although traditionally considered a zoonotic pathogen, recent genomic evidence suggests that some *C. fetus* lineages may have originated in humans before adapting to livestock during domestication (Iraola et al., 2017). The Cfv lineage includes a phenotypic variant, *C. fetus* subsp. *venerealis* biovar *intermedius* (Cfvi), which exhibits distinct biochemical traits but lacks sufficient genetic divergence to warrant classification as a separate subspecies (Iraola et al., 2013). Differentiation among these subspecies requires multilocus sequence typing (MLST), comparative genomic analysis, and host-adaptive molecular markers, as standard biochemical tests alone are often insufficient for accurate classification (Iraola et al., 2013).

Whole-genome sequencing (WGS) has become a crucial tool for exploring the genetic diversity of *Campylobacter* spp., enabling the identification of sequence types (STs), antimicrobial resistance genes (ARGs), virulence factors, and potential zoonotic transmission routes (Oniciuc et al., 2018). Previous studies have highlighted the presence of resistance to several antibiotics, including tetracycline, streptomycin, and fluoroquinolones, and they have also begun to uncover the role of virulence factors in *C. fetus* pathogenicity (Abril et al., 2010; Pena-Fernández, Ocejo, et al., 2024; Silva et al., 2021; Taylor & Chau, 1997). However, the full extent of its genomic diversity, the mechanisms underlying antimicrobial resistance (AMR), and its zoonotic potential remain poorly understood, particularly in the context of global surveillance and cross-species transmission. Although *C. fetus* harbors several ARGs (van der Graaf-van Bloois et al., 2023), comprehensive genomic studies are urgently needed to uncover the mechanisms driving AMR dissemination and the factors contributing to the persistence and spread of resistance genes. The role of mobile genetic elements (MGEs), such as plasmids and genetic islands (GIs), in *C. fetus* is an emerging area of interest, with preliminary findings indicating their potential significance in virulence, immune evasion, and AMR (Kienesberger et al., 2014).

Despite the recent advancements in understanding *C. fetus* (Pena-Fernández, Ocejo, et al., 2024; Silva et al., 2021), key questions regarding its genomic diversity, AMR, and zoonotic potential largely remain unresolved. Further more comprehensive insights into the mechanisms driving horizontal gene transfer (HGT), the role of mobile genetic elements in AMR dissemination, and the genomic features underlying host adaptation and pathogenicity are still lacking (Golz & Stingl, 2021; Kienesberger et al., 2014; Pena-Fernández et al., 2024). This study leverages publicly available *C. fetus* genomes to conduct an in-depth comparative genomic analysis, providing a global perspective on its genetic diversity, AMR patterns, and zoonotic risk across diverse hosts and geographies. By addressing these critical gaps, our findings will advance the understanding of *C. fetus* as a significant zoonotic and veterinary pathogen, offering valuable insights for global surveillance, public health strategies, and animal disease management.

## Results

### Genome data

A total of 114 *C. fetus* genomes reported high quality were retrieved from the Bacterial and Viral Bioinformatics Resource Center (BV-BRC) database, including 23 complete genomes and 91 whole-genome shotgun sequences (draft genomes). Only genomes with high completeness (>95%) and low contamination (<5%) based on CheckM estimates were included (Table S1). Excluding one outlier (strain RUG14080), genome sizes ranged from 1.7 Mb to 2.1 Mb, with GC content varying between 32.9% and 34.4%. The outlier genome, strain RUG14080, exhibited a genome size of 1.5 Mb and an unusually high GC content of 47.6%, significantly higher than the other genomes. These genomes were isolated from various hosts, with host data available for 79 genomes: the majority originated from cows (55.7%, n = 44), followed by humans (38.0%, n = 30), sheep (5.1%, n = 4), and reptiles (1.3%, n = 1). Geographical data were available for 104 genomes, with the following distribution: Europe (30.8%, n = 32), North America (24.0%, n = 25), South America (15.4%, n = 16), Asia (15.4%, n = 16), Oceania (9.6%, n = 10), and Africa (4.8%, n = 5). Further details about these genomes are provided in Supplementary Table S1.

### Taxonomic Identification and Geographic/Host Distribution of MLST Types

Taxonomic classification of the 114 *C. fetus* genomes was performed using GTDB-Tk (v2.3.2), which accurately classified all Cft genomes (n = 35) and the remaining genomes as *C. fetus* (Table S2). Subspecies assignments for all *C. fetus* genomes were further refined using secondary data (Pena-Fernández, Ocejo, et al., 2024) and validated through phylogenetic analysis. However, subspecies identification for three genomes could not be confirmed with the current analysis and available data. Of these, two genomes (SRR5279288_bin.84_CONCOCT_v1.1_MAG and first) were confirmed as *C. fetus* using GTDB-Tk, while the third genome (RUG14080) was identified as a different *Campylobacter* species (Figure S1).

A total of 32 distinct STs, ranging from ST-1 to ST-74, were identified across 111 of the 114 confirmed *C. fetus* genomes analyzed, spanning six continents and multiple host species (Table S3). The distribution varied among the three subspecies, with *C. fetus* subsp. *testudinum* (Cft), though typically associated with reptiles, being predominantly isolated from humans and exhibiting the highest ST diversity (n = 20). Overall, ST-4 was the most prevalent, representing 45 genomes, followed by ST-3, observed in 8 genomes. A substantial proportion (90.9%, n/N = 40/44) of Cfv and its biovar intermedius (Cfvi) were assigned to ST-4, with 75.6% (n/N = 34/45) originating from cattle. Europe exhibited the highest ST diversity (n = 18), while South America and Africa each recorded the lowest diversity (n = 1). In North America, ST-6 and ST-15 were the second most frequently occurring STs, each accounting for 17.4% (n/N = 4/23) of the genomes. Among genomes with host metadata, those associated with humans displayed the greatest diversity, encompassing 16 distinct STs.

### Phylogenetic and Pangenomic Analyses

The phylogenetic tree, based on core genome SNPs (1365 genes), illustrates the evolutionary relationships among *C. fetus* subspecies, revealing distinct clustering patterns that highlight genetic diversity and interrelationships (Figure 1). All Cfv and Cfvi genomes clustered closely despite being isolated from five different continents. While Cfv and Cfvi genomes formed distinct branches at the biovar level, six Cfv genomes were located within Cfvi clades, suggesting a potential shared evolutionary lineage (Figure S2). In contrast, Cft genomes, which exhibited 20 different STs, clustered together in closely associated clades, entirely distinct from the Cfv, Cfvi, and Cff groups. Human-associated genomes were primarily distributed across the Cff and Cft clades, with 60.0% (n/N = 18/30) clustering with Cft and the remaining 40.0% (n/N = 12/30) with Cff. Geographical associations were observed for Cfv and Cfvi genomes. All Cfv genomes from North America clustered on a single branch, while most Cfvi genomes from South America formed a closely related cluster. In contrast, Cff genomes displayed no clear geographical patterns, as branches often contained strains from multiple continents, indicating a more dispersed global distribution.

**Figure 1.**
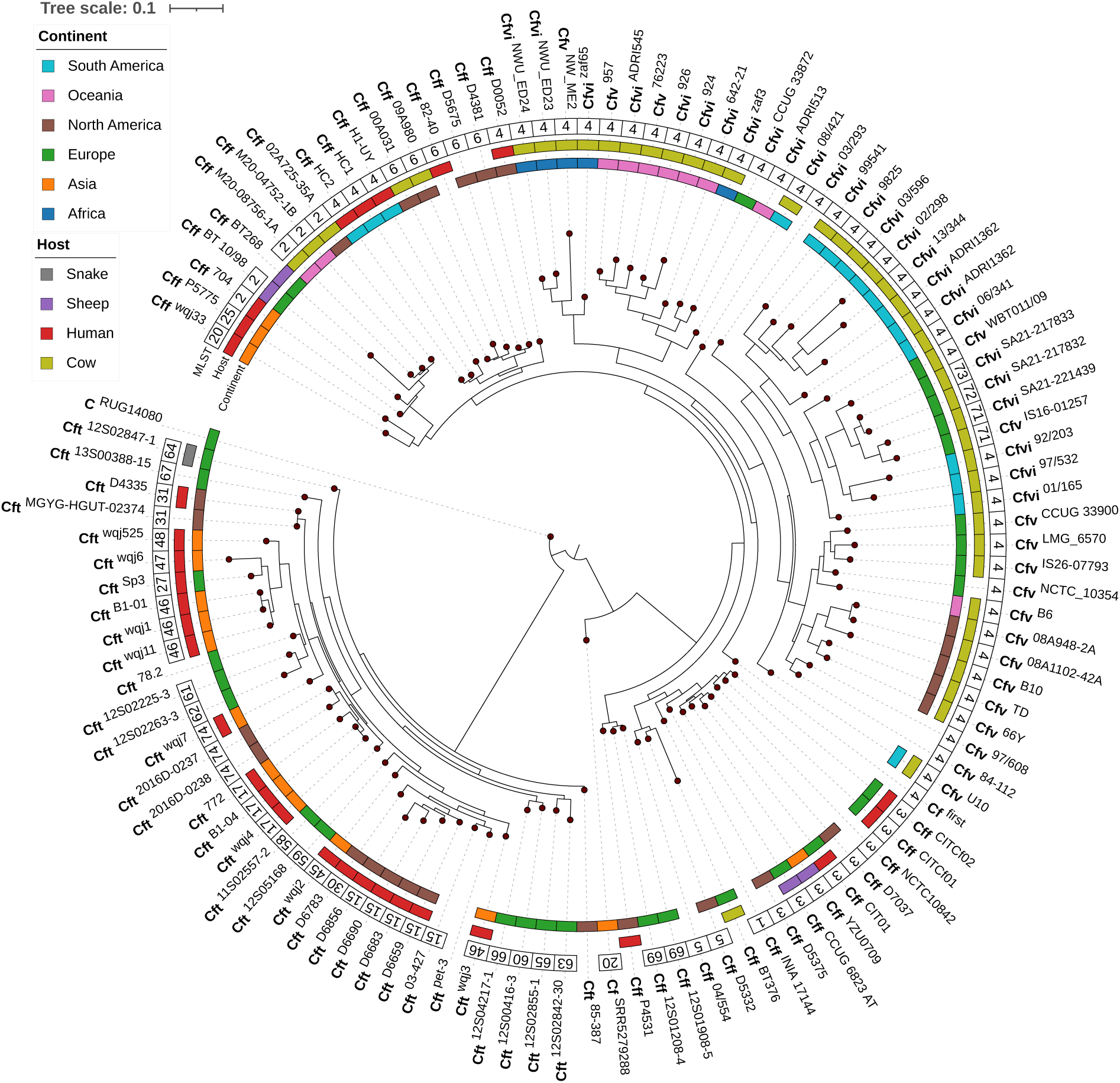
Phylogenetic tree illustrating the evolutionary relationships among the 114 *C. fetus* genomes obtained from the BV-BRC database. The tree was constructed using the Newick file generated by Roary, based on core genome SNP analysis, and visualized using the iTOL platform with metadata layers for enhanced interpretability. The inner ring represents the geographic distribution of isolates by continent, with the following colors: Africa (blue), Asia (orange), Europe (green), North America (brown), Oceania (pink), and South America (cyan). The second ring indicates host origins, represented by the following colors: Human (red), Cow (yellow-green), Sheep (purple), and Snake (gray), with blank spaces indicating genomes lacking host or geographic metadata. The third ring displays MLST data, and the outer ring depicts taxonomic classification at the species and subspecies levels (*C. fetus* [Cf], *C. fetus subsp. fetus* [Cff], *C. fetus subsp. venerealis* [Cfv], *C. fetus subsp. venerealis biovar intermedius* [Cfvi], and *C. fetus subsp. testudinum* [Cft]), with individual strain names displayed in superscript.

The analysis of 114 *C. fetus* genomes identified a total of 9,409 genes, of which 849 were core genes (469 core and 380 soft-core genes) present in at least 95% of the genomes. The remaining 8,560 genes were classified as accessory genes, consisting of 1,866 shell genes (present in 15% to 95% of strains) and 6,694 cloud genes (present in fewer than 15% of strains). The gene presence/absence matrix revealed distinct patterns among the taxonomic groups. The most pronounced clustering was observed within Cft, where a subset of genes was either unique to Cft or broadly distributed across other subspecies but largely absent in Cft. This distinct gene presence/absence pattern in the pangenome matrix resulted in the segregation of genes exclusive to Cft, as well as those absent in Cft but present in other *C. fetus* subspecies, highlighting its genomic divergence (Figure 2).

**Figure 2:**
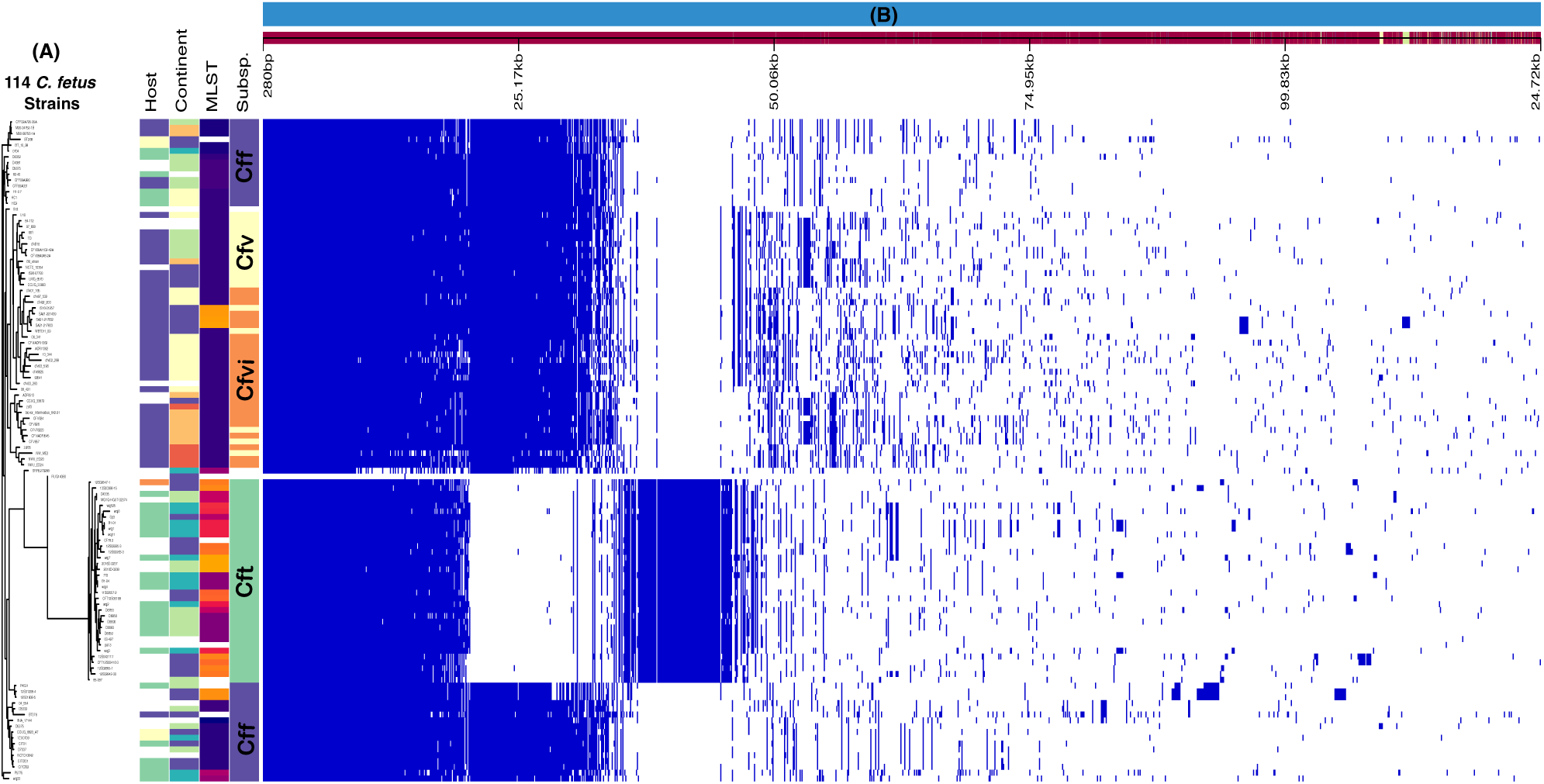
Pangenome analysis of 114 *C. fetus* genomes. (A) Dendrogram showing the clustering of 114 genomes based on accessory gene distribution, with metadata layers color-coded to indicate host species, geographic origin, MLST, and subspecies (*C. fetus* [Cf], *C. fetus* subsp. *fetus* [Cff], *C. fetus* subsp. *venerealis* [Cfv], *C. fetus* subsp. *venerealis* biovar *intermedius* [Cfvi], *C. fetus* subsp. *testudinum* [Cft]). (B) Roary matrix representing the complete genetic profile of each genome based on the presence/absence of core and accessory genes.

### Antimicrobial Resistance (AMR) Gene, virulence factors and Mobile genetic element

#### Antimicrobial Resistance Gene Profiling

AMR gene analysis across the 114 *C. fetus* genomes was conducted using Abricate, revealing the presence of five distinct AMR genes. The most commonly identified gene was *tet(O)* (n = 3), which confers resistance to tetracyclines. The distribution of these AMR genes was geographically diverse, with genes found in genomes from Asia (n = 3) and North America (n = 2) (Table 1). The two North American strains each harbored two distinct AMR genes. A comprehensive summary of the AMR genes, their genomic locations, and corresponding accession numbers is provided in Supplementary Table S4.

**Table 1.** Distribution and Frequency of Antimicrobial Resistance Genes in *C. fetus* Genomes.

| Gene | Antibiotic Resistance Conferred | Host | Region | Frequency |
| --- | --- | --- | --- | --- |
| *tet(O)* | Doxycycline, Tetracycline, Minocycline | Human | Asia, North America | 3 |
| *tet(44)* | Doxycycline, Tetracycline, Minocycline | Cow | North America | 1 |
| *ant(6)-Ib* | Streptomycin | Cow | North America | 1 |
| *aph(3')-III* | Amikacin | NA | North America | 1 |
| *lnu(C)* | Lincomycin | NA | Asia | 1 |

**Abbreviations:** NA- Non available

#### Horizontal Gene Transfer and Genomic Islands

In-depth analysis of the five genomes harboring AMR genes using Proksee revealed several HGT regions. The CARD RGI (Comprehensive Antibiotic Resistance Database (CARD) Resistance Gene Identifier (RGI)) further elucidated the resistance mechanisms associated with these AMR genes, highlighting a diverse array of resistance pathways (Figure 3). A total of 140 GIs were identified across 41 genomes, emphasizing the widespread presence of HGT elements. The distribution of genomic islands (GIs) varied across subspecies: Cff genomes contained the highest proportion (46.3%, n = 19), while Cft genomes exhibited the lowest number (12.2%, n = 5) (Table S5). Surprisingly, no plasmid sequences were detected in any of the analyzed strains using PlasmidFinder.

**Figure 3:**
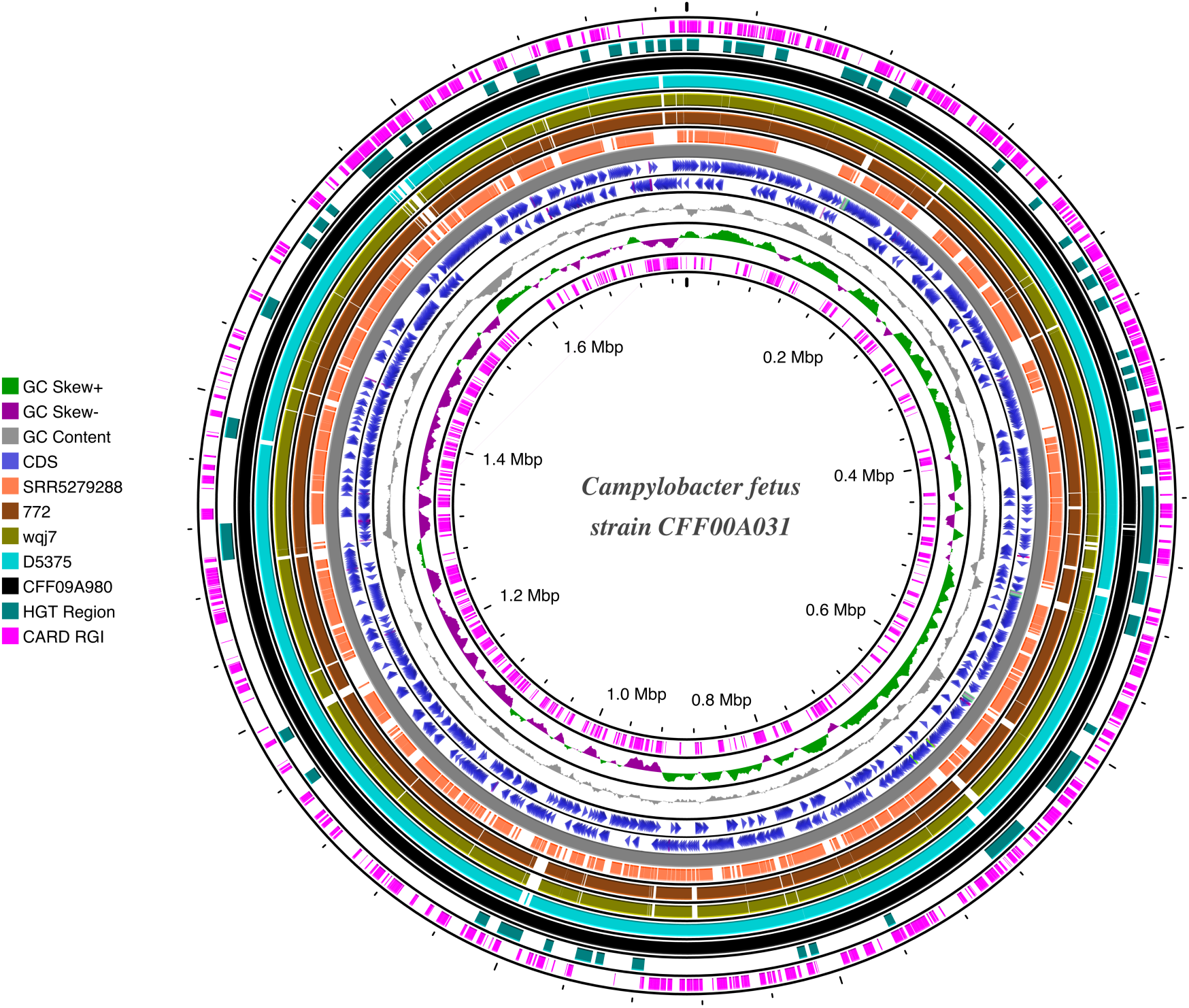
Circular genome representation of the reference genome CFF00A031 compared to five genomes harboring AMR genes. The circular visualization highlights key genomic features across multiple rings. Starting from the innermost ring: (1) predicted resistance mechanisms (pink), (2) GC skew (purple and green), (3) GC content (gray), (4 and 5) coding sequences (CDS) (two blue rings), and (6) genome backbone (solid gray). Followed by the BLASTN comparison results of the five AMR-harboring genomes in the following order: SRR5279288 (coral), 772 (brown), wqj7 (greenish-yellow), D5375 (light blue-green), and CFF09A980 (black). Horizontal gene transfer (HGT) events are indicated in the next ring marked in cyan, with the outermost ring representing the predicted AMR genes highlighted in pink.

### Virulence Factors

A virulence factor analysis of all 114 *C. fetus* genomes identified *cheY*, a key gene in the chemotaxis signaling pathway, as the sole virulence factor present. This gene was found in 35 strains across various hosts and regions (Table S6).

## Discussion

In this study, we analyzed 114 publicly available *C. fetus* genomes, confirmed their taxonomic classification and uncovered distinct geographic and host-associated patterns in ST diversity. The misclassification of strain RUG14080 at the species level, as identified by GTDB-Tk, underscores the challenges of genome-based taxonomy, particularly for strains with incomplete assemblies or unique genetic traits (Paul et al., 2019). Our findings revealed significant variation in ST distribution across continents and hosts, underscoring the global diversity of *C. fetus*. Europe exhibited the highest ST diversity, with 18 distinct STs, while South America and Africa showed the lowest diversity, each with only one ST, potentially reflecting disparities in surveillance intensity, animal husbandry practices, or pathogen prevalence (Iskandar et al., 2021; Kaakoush et al., 2015; Platts-Mills & Kosek, 2014; Sahin et al., 2017). The predominance of ST-4, particularly in cattle-associated strains (Cfv and its biovar intermedius [Cfvi]), likely reflects host-specific adaptations that enhance fitness and survival in cattle (Pena-Fernández, Ocejo, et al., 2024; van Bergen et al., 2005), coupled with clonal expansion driven by global agricultural practices such as cattle trade (Rohr et al., 2019). In North America, ST-6 and ST-15 were the second most frequently occurring STs, further highlighting regional differences in *C. fetus* populations (Kaakoush et al., 2015; Wagenaar et al., 2014). In contrast, a greater ST diversity observed in human-associated Cft strains, encompassing 16 distinct STs, highlights their zoonotic potential and warrants further investigation into interspecies transmission dynamics (Costa & Iraola, 2019). Notably, while previous studies predominantly linked Cft to reptiles (Gilbert et al., 2018), our analysis identified humans as the primary hosts for this subspecies, reinforcing its zoonotic significance.

The phylogenetic analyses of *C. fetus* subspecies provide critical insights into the evolutionary forces shaping their global diversity, host specificity, and zoonotic potential. The tight clustering of Cfv and Cfvi genomes across continents suggests the influence of conserved genomic elements that may confer adaptive advantages in host, such as immune evasion or metabolic specialization (Pena-Fernández, Ocejo, et al., 2024; Toft & Andersson, 2010). However, the placement of six Cfv genomes within Cfvi clades raises important questions regarding the potential influence of horizontal gene transfer, recombination events, or biovar misclassification (Kienesberger et al., 2014; Pena-Fernández, Ocejo, et al., 2024). This genomic overlap challenges traditional subspecies boundaries and suggests that current classification frameworks may not fully capture the complexity of genetic relationships between Cfv and Cfvi, potentially due to shared evolutionary origins and ecological adaptations (Emele et al., 2019; van der Graaf-van Bloois et al., 2020). The stark contrast in geographic clustering, with Cfv and Cfvi exhibiting regional clustering and Cff showing a more dispersed distribution, may be attributed to the host-specific nature of Cfv and Cfvi ( van der Graaf–van Bloois et al., 2016), which are geographically confined due to their association with cattle (Mshelia et al., 2008). In contrast, Cff’s broader host range and environmental resilience likely facilitate its global spread (Liu et al., 2019), potentially driven by human activities such as trade and migration. The pangenomic analysis of gene presence/absence revealed distinct patterns, particularly within Cft, where several genes absent in other subspecies were present, and vice versa, further emphasizing the unique genomic characteristics of Cft. Also, the pronounced ST diversity observed in Cft genomes underscores their broader ecological versatility, likely enabling adaptation to diverse hosts and environments (Fitzgerald et al., 2014; Gilbert et al., 2016; Patrick et al., 2013). The distribution of human-associated genomes across both Cff and Cft clades highlights the zoonotic potential of these subspecies and suggests independent evolutionary pathways facilitating interspecies transmission (Mourkas et al., 2022). These findings underscore the complex interplay of evolutionary, ecological, and anthropogenic factors shaping *C. fetus* populations and highlight the need for integrative genomic and epidemiological studies to address challenges in taxonomy, surveillance, and public health.

The AMR gene profiling across the 114 *C. fetus* genomes in this study revealed a relatively limited but geographically diverse presence of resistance genes, highlighting the potential for AMR spread within this species. The presence of AMR genes in *C. fetus* strains isolated from both human and animal hosts emphasizes the ability these strains to acquire and maintain resistance traits across different host species (Lynch et al., 2021; van der Graaf-van Bloois et al., 2023). Given the limited number of strains harboring these genes, further surveillance of *C. fetus* in diverse geographic regions and host populations is crucial to better understand the dynamics of AMR dissemination and its potential impact on public health. Nonetheless several HGT regions were identified in the analysed *C. fetus* genomes which provides further insight into the mechanisms underlying AMR gene acquisition and dissemination (Golz & Stingl, 2021). In line with the current findings an ealier study has suggested that *C. fetus* may acquire resistance genes through genetic exchange with other microorganisms (Tshipamba et al., 2020). The widespread distribution of GIs underscores the importance of HGT in shaping the genetic landscape of *C. fetus* suggesting species may be more prone to acquiring foreign genetic material, potentially enhancing its adaptability and survival in different ecological niches (Gorkiewicz et al., 2010). The absence of plasmid sequences in the analyzed strains is noteworthy, as plasmids are often key vectors for the spread of AMR genes. This raises the possibility that AMR in *C. fetus* may be predominantly mediated through chromosomal integration of resistance genes, rather than plasmid-mediated transfer (Guernier-Cambert et al., 2021). Further investigation into the role of GIs and other mobile genetic elements in the evolution of AMR in *C. fetus* is essential to unravel the full scope of genetic mechanisms that contribute to resistance in this pathogen.

Virulence factor analysis revealed that *C. fetus* strains were only haboring *cheY*, a key gene involved in the chemotaxis signaling pathway, which plays a crucial role in bacterial motility and host colonization (Foynes et al., 2000). This gene was present in 35 strains across a range of hosts and geographic regions, suggesting its potential role in facilitating *C. fetus* adaptation to diverse environments and hosts. The widespread distribution of *cheY* in both human and animal-associated strains highlights its importance in the pathogenicity of *C. fetus* and its ability to colonize different host species (Bolton, 2015; LaGier et al., 2014). However, the limited number of virulence factors identified in this study suggests that *C. fetus* may rely on other yet-to-be-identified factors for its pathogenicity, and further studies are warranted to explore the full repertoire of virulence determinants in this species. The presence of virulence factors in *C. fetus* strains from both human and animal hosts also reinforces the zoonotic potential of this pathogen, suggesting that cross-species transmission may be facilitated by the presence of conserved virulence traits (Majumdar & Pal, 2017; Miller et al., 2017). Additionally, the identification of a single virulence factor in this study contrasts with the more complex virulence profiles observed in other Campylobacter spp., highlighting the unique pathogenic strategies employed by *C. fetus*. Understanding the role of *cheY* and other potential virulence factors in *C. fetus* pathogenicity will be essential for developing targeted interventions to mitigate the impact of this pathogen on both human and animal health.

While this study provides valuable insights into the genomic diversity, AMR, and virulence potential of *C. fetus*, several limitations should be acknowledged. First, the relatively small sample size of 114 *C. fetus* genomes, though diverse in terms of geographic origin and host association, may not fully capture the genetic variability of the species across different environments or over time. The absence of plasmid sequences in the analyzed genomes, despite their role in the horizontal transfer of AMR genes, limits our understanding of the mechanisms driving AMR dissemination in *C. fetus*. Additionally, the reliance on publicly available genomic data introduces potential biases, as these genomes may not be fully representative of the entire *C. fetus* population, particularly in under-sampled regions or host species. The lack of detailed phenotypic data, such as antimicrobial susceptibility testing or in vitro virulence assays, further restricts our ability to directly correlate the presence of AMR genes and virulence factors with pathogenicity in different hosts. Finally, while subspecies classification was based on GTDB-Tk, secondary data, and phylogenetic analysis, it was not fully confirmed using more robust in silico methods, such as Kraken2, which could have provided more precise subspecies identification. Future studies incorporating larger, more diverse datasets, phenotypic data, and longitudinal sampling will be essential for a more comprehensive understanding of *C. fetus* evolution, host adaptation, and AMR mechanisms.

## Conclusion

This study provides a comprehensive genomic analysis of *C. fetus*, revealing significant genetic diversity, AMR profiles, and zoonotic potential across diverse geographic regions and host species. Our findings highlight the complexity of *C. fetus* subspecies classification, with evidence of HGT and possible subsp. misclassification, suggesting the need for more robust in silico methods for subspecies identification. While AMR genes were identified, their limited presence and the absence of plasmid sequences suggest that chromosomal integration may play a key role in resistance dissemination. Despite relying on publicly available genomes and the absence of phenotypic data, our findings emphasize the importance of integrating genomic, epidemiological, and phenotypic approaches to better understand *C. fetus* evolution, host adaptation, and AMR mechanisms, informing strategies to mitigate its impact on human and veterinary health.

## Methods

### Strain Selection

For a comprehensive genomic analysis of *C. fetus*, genomes were retrieved from the BV-BRC server (https://www.bv-brc.org/, last accessed on December 7, 2024). The search was performed using the term ‘Campylobacter fetus,’ and the filter was set to include both complete genomes and whole-genome shotgun sequences.To ensure high-quality data, only genomes flagged as good quality by the BV-BRC database were considered. Additionally, genomes were cross-referenced with NCBI taxonomy data to confirm accurate species identification (https://www.ncbi.nlm.nih.gov/datasets/genome/?taxon=196). A total of 114 *C. fetus* genomes, isolated from various hosts (cattle, humans, sheep, and reptiles) and geographically distributed across six continents, met the inclusion criteria and were retrieved for comparative genomic analysis (Table S1).

### Genome annotation and taxonomic identification

All *C. fetus* Genomes were annotated using Prokka v1.14.6 (Seemann, 2014), with default parameters. The genome annotations were outputted in GFF3 format for downstream analysis. Prokka was employed to predict genes, rRNA, tRNA, and other genomic features, and the resulting files were used for subsequent analyses. Taxonomic classification of the genomes was performed using GTDB-Tk (v2.3.2) to assign objective taxonomic classifications based on the Genome Taxonomy Database (GTDB) (Arkin et al., 2018). Subspecies assignments of Cff, Cfv, and Cfvi were further refined through phylogenetic analysis and the use of a secondary data (Pena-Fernández, Ocejo, et al., 2024).

### In-silico Multi-Locus Sequence Typing (MLST)

In-silico MLST was performed using the open-source tool MLST (https://github.com/tseemann/mlst), which queries the PubMLST database (Jolley et al., 2018). The tool identifies sequence types by aligning the genomic data to loci defined in the PubMLST database, using default parameters for allele and sequence type determination.

### Phylogenetic Analysis

Phylogenetic analysis was performed using the Newick file generated from the Roary output, which was visualized as a circular phylogenetic tree using iTOL v6.5.1 (Letunic & Bork, 2024). The tree was annotated with information on genomic features and geographical origin to investigate the evolutionary relationships between the genomes.

### Pangenome Analysis

Pangenome analysis was conducted using Roary v3.13.0 (Page et al., 2015) with a minimum sequence identity threshold of 90% for BlastP. Genes were classified into core (present in ≥99% of genomes), soft-core (95–99%), shell (15–95%), and cloud (0–15%) categories. The gene presence/absence matrix generated by Roary was visualized using the Phandango v1.3.1 web-based visualization tool (www.phandango.net) (Hadfield et al., 2018), which provided insights into the genetic diversity and evolutionary relationships within the *C. fetus* genomes.

### In silico AMR profile, virulence genes, and mobile genetic elements analysis

AMR genes and virulence factors were identified using Abricate v1.0.1 (https://github.com/tseemann/abricate) with the ResFinder (Zankari et al., 2012) and VFDB (Chen et al., 2005) databases for AMR and virulence gene prediction, respectively. Default parameters were used for gene identification, and only hits with ≥90% identity and ≥80% coverage were considered for downstream analysis. The genomes identified to be haboring AMR genes were further visualised using Proksee (https://proksee.ca/) (Grant et al., 2023). Putative HGT events were identified using Alien Hunter (Vernikos & Parkhill, 2006)

Plasmid sequences were screened for using PlasmidFinder v2.1 (Carattoli et al., 2014), implemented on the Galaxy web platform. The tool was run with default parameters, using assembled genome sequences in FASTA format as input. PlasmidFinder detects plasmid replicon sequences based on the Center for Genomic Epidemiology database

Genomic islands (GIs) were predicted using IslandPath-DIMOB v1.0.0 (https://github.com/brinkmanlab/island_path), a tool designed based on dinucleotide biases and the presence of mobility genes (Bertelli & Brinkman, 2018). The tool was run via the command line, with the default parameters and assembled genome sequences in FASTA format as input. GIs were identified based on their characteristic features, such as the presence of mobile elements and atypical dinucleotide frequencies.

## Declarations

### Ethics approval and consent to participate

Not applicable

### Consent for publication

Not applicable

### Authors’ Contributions

Conceptualization: E.K.P., L.A.O., K.O.D., and D.D.; Methodology: E.K.P., C.A.K., K.G.B., C.W.A., A.K., and S.B.; Formal analysis: E.K.P.; Data curation: C.A.K., C.W.A., and S.B.; Writing—original draft: E.K.P.; Writing—review and editing: E.K.P., L.A.O., K.O.D., D.D., C.A.K., C.W.A., K.G.B., A.K., and S.B.; Supervision: D.D., L.A.O., K.O.D., S.B., and A.K. All authors read and approved the final manuscript.

## Acknowledgements

We thank the teams and developers responsible for maintaining public genomic databases and bioinformatics tools, whose efforts were integral to this study.

## Data Availability

All data used and generated in this study are provided as supplementary information.

## Supplementary Figures

**Figure S1:**
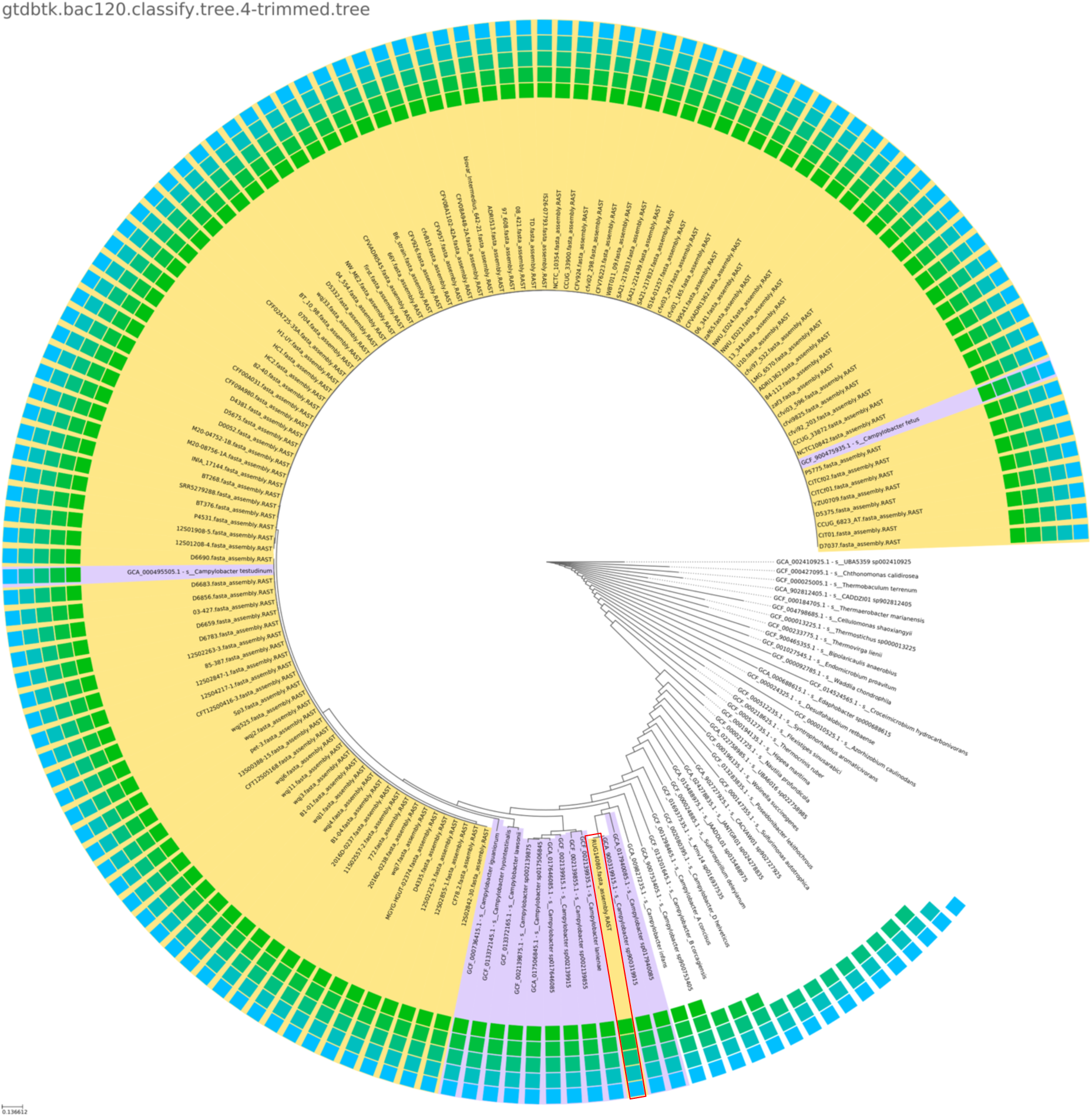
Taxonomic Identification of 114 *C. fetus* Genomes Using KBase, Highlighting Strain RUG14080 (Red) as a Different Campylobacter Species

**Figure S2:**
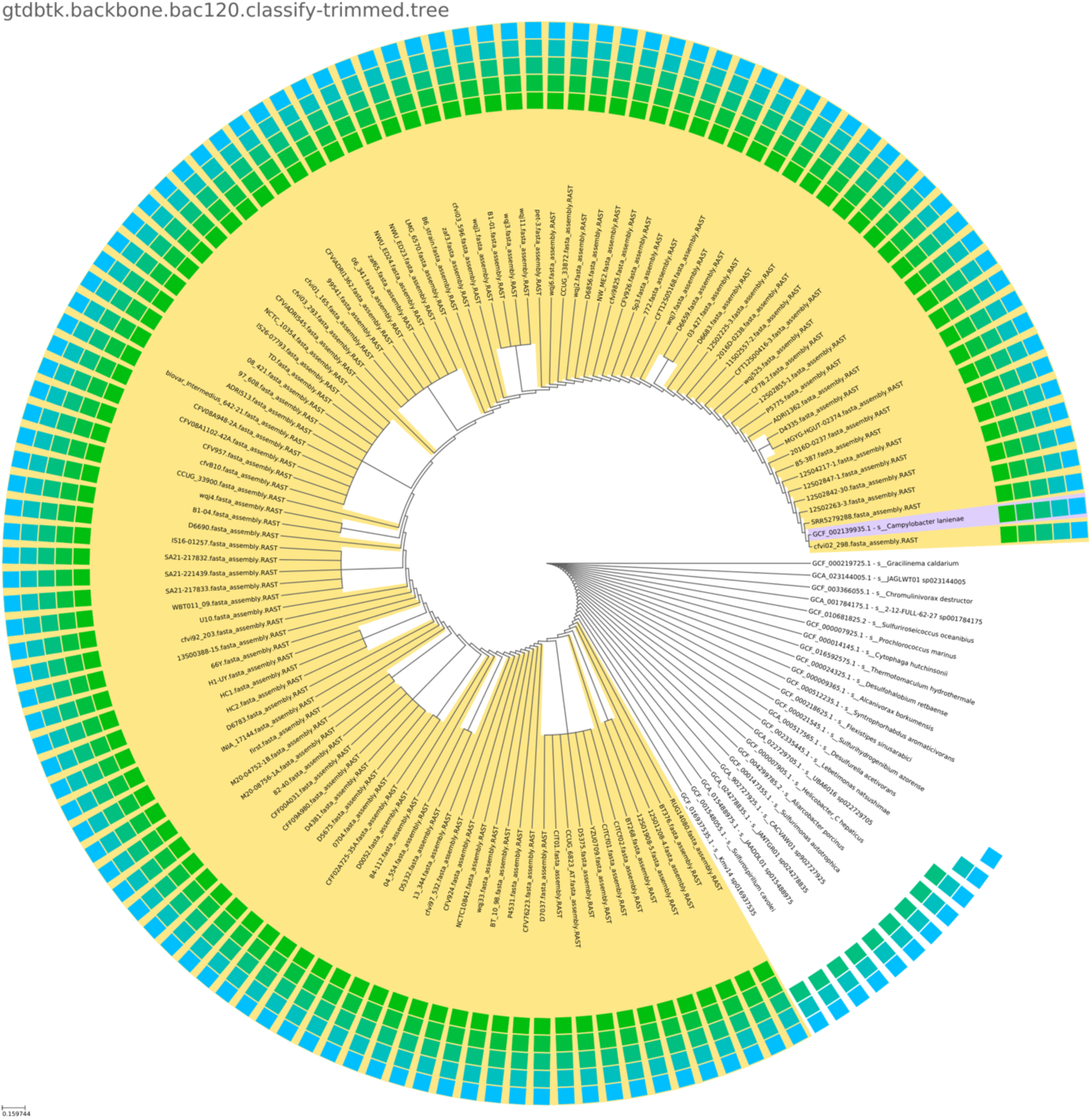
Phylogenetic tree of 114 *C. fetus* genomes generated with GTDB-Tk v2.3.2, with branch lengths indicating evolutionary distance.

